# Enhanced methods for genetic assays in *Drosophila* cells

**DOI:** 10.1101/2024.12.01.626214

**Authors:** Y Wang, JY Lee, AE Housden, E Hottinger, BE Housden

## Abstract

Genetic assays are an invaluable tool for both fundamental biological research and translational applications. Variable Dose Analysis (VDA) is an RNAi-based method for cell-based genetic assays that offers several advantages over approaches such as CRISPR and other RNAi-based methods including improved data quality (signal-to-noise ratio) and the ability to study essential genes at sub-lethal knockdown efficiency. Here we report the development of three new variants of the VDA method called high-throughput VDA (htVDA), VDA-plus and pooled-VDA. htVDA requires 10-fold reduced reagent volumes and takes advantage of liquid handling automation to allow higher throughput screens to be performed while maintaining high data quality. VDA-plus is a modified version of VDA that further improves data quality by 4.5-fold compared to standard VDA to allow highly sensitive detection of weak phenotypes. Finally, Pooled VDA allows greatly increased throughput by analysing multiple gene knockdowns in a single population of cells. Together, these new methods enhance the toolbox available for genetic assays, which will prove valuable in both high-and low-throughput applications. In particular, the low noise and ability of VDA to study essential genes at sub-lethal knockdown levels will support identification of novel drug-targets, among which essential genes are often enriched. While these tools have been developed in *Drosophila* cells, the underlying principles are transferrable to any cell culture system.

## Introduction

High-throughput genetic screens (HTS) in cultured cells have been used widely in biological and biomedical research for many years and their positive impact is clear. For example, screens in *Drosophila* cells have been used to map signalling pathways (1–3), determine mechanisms of human disease (4, 5) and investigate subcellular structures such as the nucleosome and golgi (6, 7). In mammalian cell lines, cell-based screens have been used on a massive scale to map cancer cell line dependencies as well as many other applications (8, 9). In addition, it is possible to perform genetic interaction screens in which pairwise combinations of genes are simultaneously disrupted (1, 10–13). Such screens have led to new insights into gene functions, identification of candidate anti-cancer targets as well as the mapping of molecular pathways on an unprecedented scale (8, 14, 15).

HTS experiments are limited by the need to achieve a balance between scale, without prohibitive workload or cost, and high data quality. While larger screens such as those on a genome-wide scale are often attractive due to their unbiased nature, large scale is generally linked to increased noise and limited complexity of the screen readout. Tools such as RNAi and CRISPR, as well as the protocols for their use, have been optimised over time to maximise both throughput and data quality and genome-scale screens with these tools are now relatively simple to perform (16, 17). However, screens to map genetic interactions represent a goal orders of magnitude more challenging. Genetic interaction screens on a genome scale are still far out of reach due to the large number of possible pair-wise gene combinations to be tested (approximately 200 million combinations in human cells). This problem of scale becomes even more significant due to the need for multiple independent reagents per gene and multiple replicates to reduce false-negative and false-positive results.

Screens designed to map candidate drug-targets for disease often produce inconsistent results between model systems and require extensive low-throughput follow-up work to identify the most promising candidates (18–20). There are multiple reasons underlying this issue but one contributing factor is likely to be the extent to which target genes are perturbed by different reagents or technologies. For example, CRISPR screens are often expected to generate null mutants and so produce a complete loss of function. By contrast, RNAi screens produce partial inhibition of target genes, the extent of which is dependent on the specific reagent used to target each gene. This can lead to differential results between screening methods. For example, CRISPR is expected to produce stronger phenotypes but may mask phenotypes associated with essential genes, which might be apparent with a partial knockdown of the target gene (16, 21–23). Thus, while HTS experiments have been invaluable for many research questions, the next generation of screens will require further improvements to screening methods to enhance throughput, data quality and enable greater control over gene disruption while maintaining low cost and simplicity of use.

The ability to assess partial knockdown of target genes is particularly important in synthetic lethal screens. Synthetic lethality is a type of genetic interaction in which inhibition of either of two genes individually is viable but the combined inhibition is lethal. Mapping of synthetic lethal interactions is a powerful approach for anti-cancer drug-target discovery because targeting one gene in the synthetic lethal pair, while the second gene is mutated in tumour cells is expected to specifically kill tumour cells while leaving healthy tissue unaffected (18, 19). However, synthetic lethal interactions are enriched among essential genes, therefore requiring partial knockdown of the relevant genes to enable their detection (16, 19).

To improve control over knockdown levels in HTS experiments, we previously developed a new approach to genetic loss-of-function experiments called variable dose analysis (VDA) (24, 25). This method assesses cell-based phenotypes generated from RNAi-based gene knockdown. In contrast to other approaches including dsRNA or CRISPR methods, VDA measures phenotype strength over a range of target gene knockdown efficiency in a mixed population of cells. Assessment of changes in phenotype distribution, with respect to knockdown efficiency, is measured indirectly using a fluorescent reporter. Using this approach, it is possible to assess phenotypes over a wide range of gene knockdown levels. In addition, as true phenotypes are correlated with knockdown efficiency, this approach allows efficient separation of signal from noise, which primarily affects the strength of phenotype rather than the phenotype distribution. VDA therefore results in improved signal-to-noise ratio compared to dsRNA assays (24). We have applied this approach to synthetic lethal screens in *Drosophila* cells and identified several candidate drug-targets for tuberous sclerosis complex and neurofibromatosis type 1, which were validated in multiple other cell culture and *in vivo* models (24, 26–28).

Here, we report new methods based on VDA that address some of the remaining challenges in HTS experiments. We have developed three variants of the approach, which can be applied to distinct experimental needs. First, we established a pseudo-miniaturised version of VDA called high-throughput VDA (htVDA). This method combines miniaturised liquid handling automation, a variant transfection technique specific to VDA and an optimised shRNA expression vector to reduce reagent usage by up to 10-fold, with no significant effect on data quality. Second, the VDA-plus method combines improvements to the shRNA expression vector with machine learning-based data analysis to greatly increase signal-to-noise ratio compared to standard VDA, resulting in a highly sensitive assay. Finally, pooled VDA takes advantage of the increased signal-to-noise ratio of VDA-plus to simultaneously assess multiple shRNAs in a single sample, greatly improving throughput. Together, these tools enable the application of VDA with improved efficiency and quality to both low and high-throughput applications.

## Results

### Optimisation of the VDA reagent delivery method

Our previously published protocols for VDA assays require the use of lipid-based transfection to simultaneously deliver multiple plasmid-encoded components into cultured cells (24, 25). Specifically, three plasmids are co-transfected, encoding a Gal4 protein, shRNA construct (the expression of which is controlled by Gal4 using the UAS system (29)) and a GFP expressing plasmid to report relative transfection efficiency per cell and therefore target-gene knockdown efficiency on a single cell level. It is standard practice when using transfection to optimise conditions to maximise delivery of plasmids to the cell population. This is generally measured based on the total signal from a reporter gene transfected into a cell population or by the proportion of cells receiving the plasmid. However, the transfection requirements for VDA are different from most other applications of transfection. VDA requires a wide range of plasmid doses to be delivered across individual cells in the population but does not necessarily require the total plasmid dose to be high. In addition, the proportion of cells transfected is not critical because the presence of the fluorescent reporter enables identification of transfected cells during data acquisition. Given these specific requirements, it is important to optimise transfection conditions to achieve the best performance from VDA assays (**File S1**).

### Transfection dose and Gal4 expression contribute to non-specific toxicity

First, we sought to minimise non-specific toxicity associated with the transfection process. Non-specific cell toxicity can be induced when cell populations are transfected with a high dose of transfection reagents. In addition, expression of transgenes encoded by the transfected plasmids can also lead to toxicity that is not due to knockdown of the target gene. To assess the effects of transfection dose, we transfected cells with varying volume of transfection mix containing a GFP expression plasmid and used flow cytometry to simultaneously assess cell viability (measured using forward scatter and side scatter; **Figure S1**), transfection efficiency (percentage of cells transfected) and transfection dose (relative average amount of plasmid delivered per cell) in wild-type *Drosophila* S2R+ cells. Cells were transfected with 1 µl, 2 µl, 3 µl or 5 µl of transfection mix. As anticipated, increasing transfection complex volume was associated with reduced cell viability, increased transfection efficiency and increased plasmid dose per cell (**Figure 1A**). As described above, reduction of the transfection efficiency or dose may not represent a significant problem for VDA assays. Therefore, it may be possible to reduce toxicity by decreasing the transfection volume without adversely affecting data quality.

**Figure 1:**
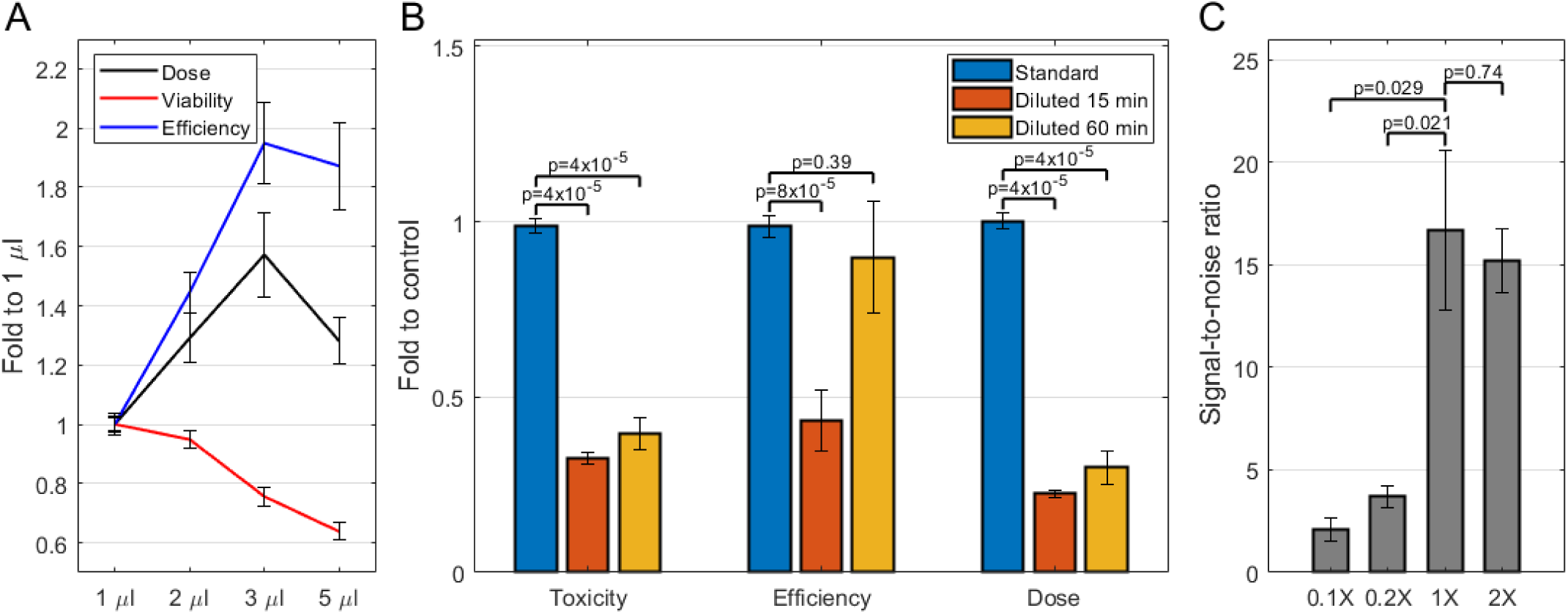
Optimisation of the VDA method. **A:** Graph showing the fold change in plasmid dose (average GFP signal per cell), viability (based on FSC/SSC gating) and transfection efficiency (proportion of cells expressing GFP) resulting from transfection with different volumes of transfection complex. Each point represents the average of at least 20 biological replicates and error bars indicate standard error of the mean. **B:** Graph showing the fold change in toxicity (measured using propidium iodide staining), transfection efficiency and plasmid dose compared to the unmodified transfection protocol. ‘Diluted’ indicates that a 10-fold dilution of transfection reagents was used and ‘15 min’ or ‘60 min’ indicates the incubation time for formation of transfection complexes. Each bar indicates the average of 9 biological replicates and error bars indicate standard error of the mean. P-values were calculated using a Wilcoxon rank sum test **C:** Graph indicates signal-to noise ratio for VDA assays performed using various dilutions of transfection reagents as indicated. Each bar represents the average of three signal-to-noise ratio values, each generated from a group of approximately 30 biological replicates. Error bars indicate standard error of the mean between the three groups of replicates. P-values were calculated using unpaired, two-tailed t-tests

Next, we assessed whether any of the plasmids used in VDA assays were associated with toxicity. We assessed a range of transfection conditions; varying the total amount of Gal4 plasmid, shRNA plasmid or combined Gal4 and shRNA plasmid delivered as well as the ratio between the two plasmids, using forward scatter (FSC) and side scatter (SSC) as a rapid means to assess cell viability (**Figure S2A-F)**. We found that variation in most of these variables had little effect on cell viability. However, high dose of the Gal4 plasmid was associated with a reduction in cell viability (**Figures S2A and S2E**), suggesting that the use of this expression system may not be optimal for VDA assays.

These results suggest that the quality of data obtained from VDA assays may be improved by reducing background toxicity through the use of a reduced volume of transfection reagent and by changing the expression system to avoid the use of Gal4.

### Pseudo-miniaturisation of VDA assays

The observation above that the use of low transection volume may enable low toxicity without significantly affecting data quality from VDA assays is consistent with our previously published VDA protocol, where we show that addition of only 1 µl transfection complex is sufficient to generate robust signals from VDA assays (25). To enable simple manual pipetting, it is necessary to generate larger volumes of transfection mix and we generally produce 10 µl for use in a single well of a 96-well cell culture microplate. This 10-fold reduction in transfection complex volume necessitates pipetting very low volumes of transfection complex components (e.g. 60 nl of transfection reagent per sample). We therefore employed a Mosquito LV Genomics low-volume liquid handling robot to assess the performance of transfections miniaturised by 10-fold. Our initial attempts to miniaturise transfection complex generation were unsuccessful due to the rapid evaporation of components during pipetting. Similarly, while the use of mineral oil was effective at preventing evaporation (**Figure S3**), transfection efficiency was severely reduced, suggesting that transfection complex formation is incompatible with the presence of mineral oil.

Given the difficulties encountered with miniaturising the transfections, we instead investigated the possibility of generating transfection complexes at reduced reagent concentrations; while maintaining the total volume at 10 µl. We tested whether decreasing the concentration of DNA and transfection reagent by 10-fold would adversely affect transfection. We found that toxicity, transfection efficiency and plasmid dose per cell were all significantly reduced (**Figure 1B**, grey bars). We hypothesised that the reduction in reagent concentration would reduce the rate of transfection complex formation but would not necessarily alter the process per se. We therefore tested whether increased incubation time would allow improved complex formation and enhance transfection efficiency. Consistent with our hypothesis, increased incubation time (from 15 to 60 minutes) resulted in improved transfection efficiency similar to levels achieved using the standard method and no significant increase in toxicity. However, plasmid dose remained relatively low (**Figure 1B**, orange bars).

VDA assays measure the relationship between phenotype and target gene knockdown efficiency. Therefore, while it is necessary to generate a range of plasmid doses in order to produce variable knockdown efficiency, the absolute plasmid dose itself is not necessarily critical. To determine whether the reduced plasmid dose would affect VDA assay output, we calculated signal-to-noise ratios (difference between means divided by the sum of standard deviations for positive and negative controls) for transfections performed with 10-fold (0.1X) and 5-fold (0.2X) reduced reagent concentrations as well as control conditions (1X) and two-fold increased concentration (2X). For conditions with reduced reagent concentrations, incubation times were increased from 15 to 60 minutes. Reducing reagent concentration resulted in a significant reduction in signal-to-noise ratio (8.0-fold for 0.1X and 4.5-fold for 0.2X) in both cases compared to 1X control (**Figure 1C**). However, increased concentration did not result in significantly improved signal-to-noise ratio.

A key requirement of VDA assays is that delivery of the GFP reporter plasmid correlates with delivery of the shRNA expression plasmid. Given that dilution of the transfection mixture results in a reduction in total plasmid dose delivered to each cell, but a similar transfection efficiency compared to the standard method, it is possible that the transfection complexes formed are smaller (i.e. contain less plasmid DNA) using this method, resulting in increased variation in the proportion of each plasmid present in each complex. We therefore hypothesised that the reduction in signal-to-noise ratio was caused by a loss of correlation between RNAi expression and GFP reporter expression. To assess this possibility, we tested whether reagent dilution affected the co-delivery of two plasmids expressing GFP or mCherry fluorescent proteins. Using the standard transfection protocol, the two fluorophores were well-correlated but this correlation was significantly reduced when transfections were performed using the diluted transfection protocol (**Figure 2A-C**). These data suggest that the loss of correlation between shRNA expression and GFP reporter expression is likely to be responsible for the reduced signal-to-noise ratio. This effect is likely exacerbated by the need to deliver three correlated plasmids (encoding shRNA, Gal4 and GFP) for VDA assays to function.

**Figure 2:**
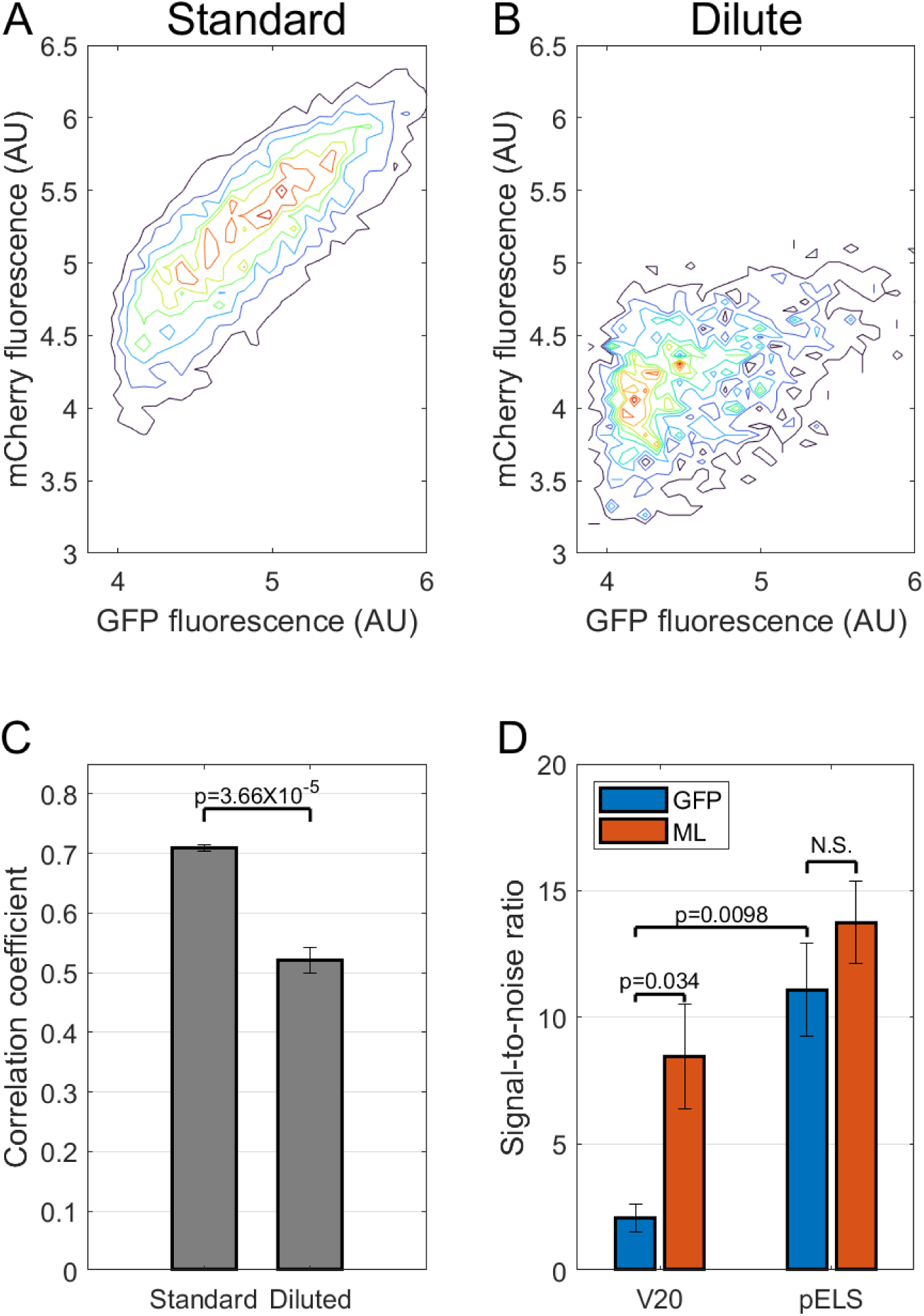
Dilution of transfection complexes is an effective way to reduce reagent use in VDA assays. **A-B:** Density plots illustrate the distribution of signals from co-transfected plasmids using a standard transfection method (A) or the diluted method (B). **C:** Quantification of correlation between the signals from co-transfected plasmids. Bars represent the average of 12 biological replicates and error bars indicate standard error of the mean. The p-value was generated using a Wilcoxon rank sum test. **D:** Graph indicates signal-to noise ratio for VDA assays performed using the pValium20 or pELS vector, and analysed using the standard VDA method or the machine learning method. Each bar represents the average of three signal-to-noise ratio values, each generated from a group of approximately 30 biological replicates. Error bars indicate standard error of the mean between the three groups of replicates. P-values were generated using unpaired, two-tailed, t-tests. N.S. indicates p>0.05.

### Signal-to-noise ratio is enhanced using a new expression system

To overcome this issue, we generated a new vector (pELS) that drives shRNA expression using a constitutively active *Act5c* promoter (**Figure S4**). The use of this promoter avoids the need to supply Gal4 and therefore reduces the number of plasmids that must be co- ransfected while also removing the non-specific toxicity associated with the Gal4 plasmid (**Figure S2A**). Measurement of signal-to-noise ratio using the pELS vector to express shRNA targeting *white* or *thread* showed a considerable increase in data quality from 2 to 11 signal-to-noise ratio using the diluted transfection method (**Figure 2D**). This is likely driven by improved co-delivery of plasmids as well as reduced toxicity caused by expression of Gal4.

### Machine learning-based data analysis enhances signal-to-noise ratio using existing reagent libraries

While the pELS vector enables the use of VDA assays with 10-fold reduced reagent volume, while maintaining high data quality, it requires the generation of new shRNA expression constructs. Reagent libraries have already been generated targeting 6,644 genes (47% of the *Drosophila* genome) using the pV20 vector (30, 31). To enable compatibility between the diluted VDA method and these existing libraries, we explored an alternative method to overcome the issues associated with loss of correlated plasmid delivery.

Transfection of cells with shRNAs targeting essential genes (e.g. *thread*) causes alterations in light scattering properties of the cells, affecting both the forward scatter (FSC) and side scatter (SSC) signals measured during flow cytometry analysis (**Figure 3A-B**). We therefore considered whether analysis of a combination of GFP, FSC and SSC measurements would result in more reliable identification of cell viability phenotypes. To achieve this, we applied machine learning. We performed VDA assays using either *white* or *thread* shRNA and using the pELS vector, with either 2X or 0.1X dilution of transfection reagents. We then selected one third of the data and used this to train a series of standard machine learning algorithms. Each cell from the training data was considered individually and assigned as having originated from a reduced viability (transfected with *thread* shRNA) population or a control population (transfected with *white* shRNA). 25% of the data were reserved as a holdout to assess performance of each algorithm and prevent over-fitting. In addition, we assessed the speed of the resulting trained algorithms to enable high-throughput data analysis (**Table 1**). While several of the machine learning methods resulted in high prediction accuracy, there was high variation in analysis speed. Based on these two parameters, we selected the Fine Decision Tree (FDT) model for further analysis, which showed the highest prediction accuracy and among the greatest prediction speeds.

**Figure 3:**
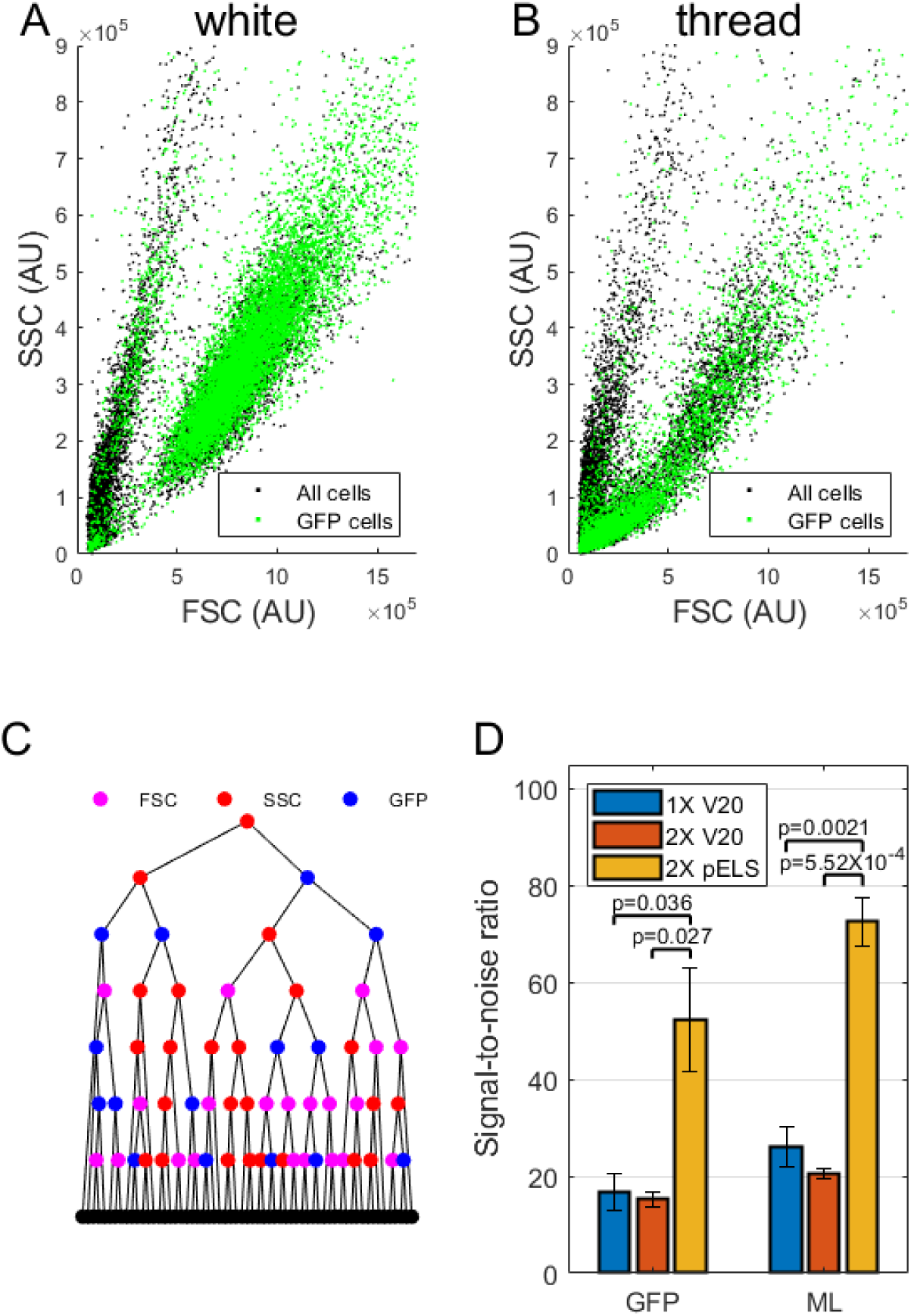
Machine learning improves data quality from diluted VDA assays. **A-B:** Scatter plots illustrate the effect of reduced cell viability on the FSC and SSC readings from individual cells. Each point represents a single cell from a population transfected with a negative control (white shRNA; A) or with a reagent to reduce cell viability (thread shRNA; B). **C:** Schematic illustrating the process and data types used to predict cell viability phenotypes by the Fine Decision Tree algorithm. **D:** Signal-to-noise ratios associated with the use of the pValium20 or pELS vector and comparing analysis methods. Bars represent the average of three signal-to-noise ratio values, each generated from a group of approximately 30 biological replicates. Error bars indicate standard error of the mean between the three groups of replicates. P-values were calculated using unpaired, two-tailed t-tests.

**Table 1:** Summary results from 23 machine learning algorithms.

| ML Algorithm | Validation Accuracy (%) | Prediction Speed (obs/s) | Interpretability |
| --- | --- | --- | --- |
| Fine Decision Tree | 81.9 | 1,400,000 | Easy |
| Medium Decision Tree | 81.5 | 1,400,000 | Easy |
| Coarse Decision Tree | 81.2 | 1,300,000 | Easy |
| Linear Discriminant | 78.2 | 880,000 | Easy |
| Quadratic Discriminant | 57.1 | 870,000 | Easy |
| Logistic Regression | 78.3 | 1,300,000 | Easy |
| Linear SVM | 54.6 | 710 | Easy |
| Quadratic SVM | 56.9 | 46,000 | Hard |
| Cubic SVM | 57.0 | 130,000 | Hard |
| Fine Gaussian SVM | 82.9 | 220 | Hard |
| Medium Gaussian SVM | 81.9 | 190 | Hard |
| Coarse Gaussian SVM | 81.1 | 170 | Hard |
| Fine KNN | 76.5 | 32,000 | Hard |
| Medium KNN | 82.4 | 14,000 | Hard |
| Coarse KNN | 82.5 | 4,500 | Hard |
| Cosine KNN | 81.5 | 120 | Hard |
| Cubic KNN | 82.5 | 3,700 | Hard |
| Weighted KNN | 81.9 | 13,000 | Hard |
| Boosted Trees Ensemble | 81.5 | 70,000 | Hard |
| Bagged Trees Ensemble | 83.1 | 19,000 | Hard |
| Subspace Discriminant Ensemble | 79.3 | 48,000 | Hard |
| Subspace KNN Ensemble | 81.4 | 4,700 | Hard |
| RUSBoosted Trees Ensemble | 81.4 | 95,000 | Hard |

To determine whether the algorithm was transferrable to other datasets, we first applied it to the two-thirds of the data not previously used for training and calculated signal-to-noise ratios for each dataset. Estimates of VDA signal were calculated by taking the sum of certainty scores generated by the FDT model over each population of cells. Results were similar to those produced from the training dataset, indicating that the model is transferrable between datasets and is not significantly over fitted. To assess how the tree model was classifying cells, we analysed which aspects of the data were used (**Figure 3C**). The tree contains 61 different decision routes. Of these, 56 use all three types of data (FSC, SSC and GFP) and 5 use only SSC and GFP. This indicates that the most powerful method for detecting cells with reduced viability is to combine all three data types together.

To further assess the machine learning analysis algorithm, we applied it to data generated using both the pV20 and pELS vectors as well as multiple transfection methods. In all cases, the machine learning algorithm produced higher signal-to-noise ratio compared to any other analysis method (**Figure 2D** and **Figure 3D**). When used with V20 plasmids and the 10-fold diluted transfection method, the ML analysis resulted in a signal-to-noise ratio of approximately 8 (**Figure 2D**), which is sufficient for HTS applications in which relatively robust phenotypes are expected. Thus, diluted transfections can be applied to screens employing existing libraries of V20 shRNA reagents to reduce costs and increase screen scale.

In summary, by altering the shRNA expression system to reduce the need for co-delivery of multiple plasmids and applying machine-learning-based data analysis, it is possible to reduce reagent use by 10-fold in VDA assays while maintaining high data quality. This will allow larger screens to be performed without major cost implications. Such advances are important for the economy of HTS experiments in general but also in the progression towards the scale of screens required to map genetic interactions on a large scale. We named this variant of the VDA method high-throughput VDA (htVDA).

### VDA-plus is a high-sensitivity version of the VDA method

Having developed improved methods for shRNA expression and data analysis that allow htVDA assays to be performed while maintaining high data quality, we tested whether the application of these improvements to standard VDA assays without dilution of transfection complexes would further enhance data quality. We performed VDA assays with 1x or 2x reagent concentration using the pV20 or pELS vector to express positive (*thread*) or negative (*white*) control shRNA reagents. We then analysed effects using either the standard GFP-based method or the ML method. Consistent with previous results (**Figure 1C**), increasing the reagent concentration did not have any significant effect on signal-to-noise ratio (**Figure 3D**, orange and grey bars). This is likely because correlation between co-transfected plasmids is already maximal under these conditions. Conversely, the use of the pELS vector had a large effect on signal-to-noise ratio and the ML-based analysis method produced a further increase in data quality although this increase did not meet the threshold for statistical significance (**Figure 3D**, purple bars). Together, the application of the new vector and new analysis method resulted in signal-to-noise ratio of approximately 72, representing a 4.5-fold increase compared to the previous VDA method and a 12-fold increase over our previous estimates of signal-to-noise ratio for dsRNA-based assays performed under similar conditions (24). VDA- plus therefore represents a high sensitivity approach ideal for analysis of weak or variable phenotypes.

### Development of a partially-pooled VDA method to increase throughput

The choice of assay in HTS often involves a trade-off between throughput (with associated cost) and data quality. Having optimised the signal-to-noise ratio of VDA assays, we next explored methods to increase assay throughput by pooling reagents into a single population of cells.

We previously showed that VDA assays could reliably detect cell viability phenotypes even when reagents were diluted up to 250-fold (24). Given the further improvements in signal-to-noise ratio achieved with the VDA-plus assay, it is likely that phenotypes could be reliably detected even when shRNA reagents are present at very low doses. For example, it may be possible to detect phenotypes from a single shRNA present in pooled assays in which multiple shRNA reagents are mixed, with each being present at greatly reduced dose compared to individual assays.

To test this, we first generated a pooled library of 45 shRNA reagents targeting a selection of kinases in the pELS vector (**Table S1, KL1 library**). We then added either a positive control shRNA (targeting *thread*) or a negative control shRNA (targeting *white*) to the library and compared results from subsequent VDA assays. We found that significant differences in phenotype could be reliably detected using these pooled assays despite the 46-fold dilution of the control reagents (**Figure 4A**).

**Figure 4:**
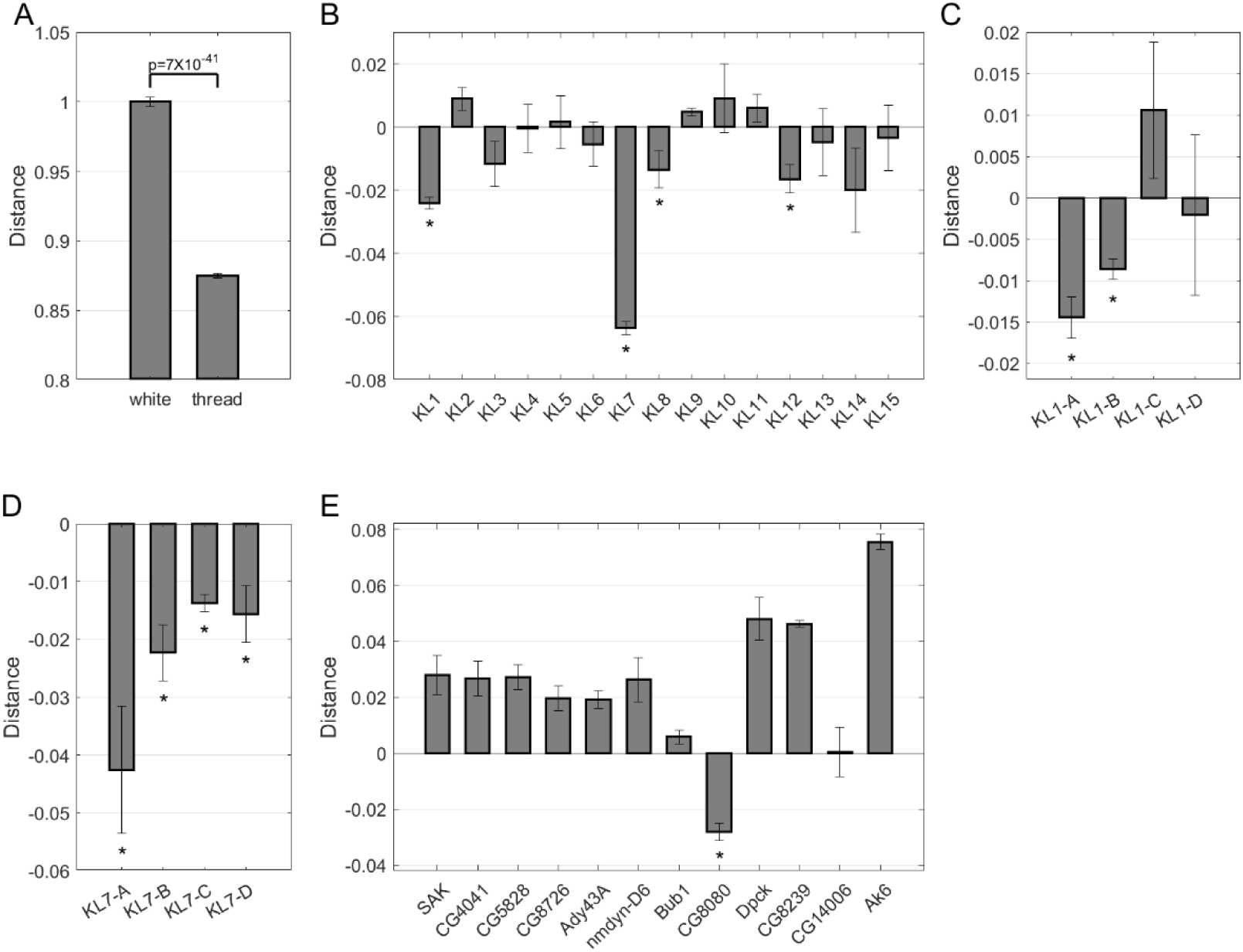
Pooled-VDA is an effective method for detection of synthetic lethal interactions. **A-E:** All graphs indicate the deviation in ratio of VDA signal between wild-type cells and NF1-deficient cells from the expected value for the shRNA reagents, or pools of reagents, indicated (see Materials and Methods for details of the analysis). Bars indicate the average from at least three biological replicates and error bars indicate standard error of the mean. P-values were calculated using z-tests and asterisks indicate p<0.05.

Next, we further tested the method by generating fifteen pooled libraries each containing up to 48 shRNA reagents targeting kinases. In total, this library includes 675 reagents covering all kinases in the *Drosophila* genome with the majority of genes targeted with two reagents (**Table S1**). We then used these libraries to screen for synthetic lethal interactions with the tumour suppressor gene *NF1*. We have previously performed genome-wide synthetic lethal screens for *NF1* (28) and so could compare results from these pooled assays to a previously validated dataset. Only one kinase was identified as synthetic lethal with *NF1* in the previous genome-wide screens. This could therefore be used as a positive control to validate the pooled-VDA assay.

Results from the assays showed that libraries KL1, KL7, KL8 and KL12 had a significantly greater effect on the viability of NF1 cells compared to wild-type control cells, suggesting a synthetic lethal interaction (**Figure 4B**). We focused on the KL1 and KL7 libraries as these produced the strongest phenotypes. The KL1 pool contained a shRNA targeting the *ND-42* gene, the positive control interactor that was previous identified in genome-wide synthetic lethal screens. No known synthetic lethal interactors were represented by the KL7 pool, suggesting that this result may indicate a novel interaction. To identify the responsible gene in each pool, we generated sub-libraries containing 11-13 shRNAs each representing subsets of the original pools. Sub-library KL1-A Produced the strongest synthetic lethal effect, consistent with the presence of the positive control shRNA targeting *ND-42* in that sub-library (**Figure 4C**). Sub-library KL7-A produced the strongest synthetic lethal effect compared to other sub-libraries from the KL7 pool (**Figure 4D**). We therefore tested the 12 shRNAs present in that sub-library individually to identify the responsible reagent. We found that only one shRNA produced a synthetic lethal effect with *NF1* when tested alone, *CG8080* (**Figure 4E**). This gene encodes a conserved mitochondrial NAD kinase (NADK2 in humans). A second shRNA reagent targeting *CG8080* was present in the KL14 library, which produced the third strongest selective viability effect on NF1 cells in the initial screen, although the effect did not meet the threshold for statistical significance (p=0.066) (**Figure 4B**).

In summary, our efforts to optimise the VDA method have resulted in a tuneable system capable of delivering high quality data or data comparable to the previous methods but with increased throughput. These new methods will increase the quality and economy of future screens performed using the VDA method.

## Discussion

The technologies available for HTS experiments have developed greatly over the past 20 years (16, 17). Because of this, it is now possible to perform genome-wide loss-of-function analysis using RNAi or CRISPR screens across a wide range of different cell-based model systems. However, despite these advances, there are still limitations in our ability to perturb gene function in a controlled manner at high throughput. For example, while we can delete genes or reduce their expression using CRISPR or RNAi, more nuanced control of expression levels remains challenging. Methods such as RNAi or CRISPRi enable different levels of gene expression to be assessed but this generally requires the use of multiple reagents, each inhibiting gene function to a different extent. VDA enables the effects of many different levels of gene expression to be assessed in a single population of cells (24, 25). However, the original version of VDA was a relatively slow and relatively expensive process, limiting the scale at which experiments could be performed.

Here, we have establish new variants of the VDA method, designed to overcome some of the previous limitations. We provide multiple methodological options, enabling the most appropriate method to be chosen for the relevant biological question and experimental limitations. First, htVDA allows arrayed screens to be performed with 10-fold reduced reagent costs, allowing larger screens but without a significant reduction in signal-to-noise ratio compared to the original VDA method. This approach can be performed using optimised vectors to express shRNA or using existing reagents in combination with machine learning-based data analysis. Existing reagents include the >6,500 shRNA reagents available in the TRiP library (31). This backward compatibility avoids the need to generate new libraries for large scale screening with this method. We note that the htVDA method involves reduced plasmid doses being delivered compared to other VDA methods. It is possible that this will result in detection of different sub-classes of phenotypes. For example, phenotypes that only occur with high efficiency knockdown may be less easily detected with this approach.

Second, VDA-plus takes advantage of some of the same developments used in the htVDA method but trades throughput for a considerable increase in signal-to-noise ratio. This allows highly sensitive assays to be performed, which will allow detection of weak phenotypes or the use of noisy assays, which otherwise would not be suitable for HTS experiments. This may be particularly useful in drug-target screening applications where the choice of assay may be limited by the nature of the disease under consideration.

Furthermore, the increase in signal-to-noise ratio associated with VDA-plus assays mean that reagents can be pooled and assessed in combination using the pooled-VDA method. While this results in dilution of the signal from any individual reagent, the high sensitivity of the assay makes it possible to detect phenotypes despite this dilution, thereby allowing much more rapid screening of reagents. We demonstrate the value of this by screening a library of 675 shRNA reagents in only 15 cell populations. This pooled-VDA method will therefore allow much larger screens to be performed in the future including genome-wide genetic screens. VDA has the added advantage of being able to assess phenotypes over a range of knockdown efficiencies, which no other technology can currently achieve at scale. This will be especially important for the identification of phenotypes associated with essential genes, which has proven challenging in previous HTS projects. In particular, synthetic lethal interactions are highly enriched among essential genes (19, 24, 32) and so drug-target screens targeting these interactions are likely to benefit greatly from these advances.

The next frontier in functional genomics is likely to be large scale mapping of genetic interactions using high-throughput screens. Pioneering studies in yeast have demonstrated the power of such screens to elucidate the functions of individual genes, map pathway structures and identify candidate drug targets for disease (10, 14, 33, 34). Genetic interaction screens have also been performed on relatively small subsets of genes in *Drosophila* and mammalian cells, demonstrating similarly powerful outputs compared to the yeast screens (1, 15, 35, 36). However, current technologies are not sufficiently scalable to advance these studies to genome-scale in these more complex model systems. The pooled-VDA method offers a potential solution to this limitation. Most genetic interaction screening methods rely on maintaining separation of reagents into different cells to allow accurate attribution of phenotypes to reagents. However, pooled-VDA allows many reagents to be present in the same cells and simply reads the average signal between those reagents with sufficient sensitivity to detect diluted phenotypes. This averaged phenotype include all single reagent effects but also all effects resulting from combinations of reagents present in the cells. Genetic interactions could therefore be identified using a similar approach to that used for the single-gene effects where sub-libraries are used to narrow down the list of potentially causative reagents. While we have not tested this possibility here, it represents a potentially powerful opportunity for future development to enable large-scale genetic interaction screens.

Finally, while VDA offers various advantages over CRISPR and other RNAi-based methods for loss-of-function assays, it has so far only been applied in *Drosophila* cells. In principle, we expect the method to be effective in any cell line that can be transfected with standard protocols. The establishment of protocols for the application of VDA assays in a wider range of cell culture systems would likely be transformational to HTS, applied to a broad range of applications. For example, to enable efficient drug-target screens in human cells for diseases that cannot be readily modelled in *Drosophila* cell culture systems. Similarly, the development of further modifications to the VDA method to enable a broader range of readouts would be valuable. For example, the use of imaging cytometry would allow the use of high-content imaging readouts while maintaining the ability to rapidly measure single-cell knockdown efficiencies.

## Methods

### Cell culture

*Drosophila* S2R+ cells were cultured using Schneider’s medium with 10% FBS and 1% penicillin/streptomycin. Cells were maintained at 25°C in T25 flasks and passaged approximately twice weekly. Cells were confirmed mycoplasma free through regular testing.

### Cell viability, toxicity and transfection efficiency assays

Forward scatter (FSC) and side scatter (SSC) were used as a rapid measurement of cell viability for initial analysis of transfection methods. Cytometry data were gated as illustrated in **Figure S1A** and the percentage of events within the P1 gate used to determine viability.

Cell toxicity was measured using propidium iodide (PI) staining. PI was added to the cell suspension at a concentration of 1:100. Samples were incubated for 5 minutes before flow cytometry analysis. PI signal was detected using the ECD channel (Excitation: 561 nm, Emission bandpass: 610/20). The percentage of transfected cells positive for PI staining was determined using the gating strategy illustrated in **Figure S1D-G**. First a gate was used to remove noise due to cell debris and other small objects in the sample using FSC and SSC (**Figure S1D**). Next, doublets were removed using a FSC-H and FSC-A gate (**Figure S1E**). Transfected cells were identified using FITC signal (**Figure S1F**) and finally, PI positive cells were identified using ECD signal (**Figure S1G**).

Transfection efficiency (proportion of cells transfected) was measured using the gating strategy shown in **Figure S1A-C**, with the percentage of events in gate P3 used to determine efficiency. Transfection dose was determined by calculating the average FITC between events within this gate.

### Machine learning training and analysis

Training of machine learning algorithms was performed using the MATLAB Classification Learner app. For training, data from 30 replicate transfections with each of the *white* and *thread* shRNA reagents using the standard transfection method (with the pELS vector) and 30 additional replicates using the same reagents but with the 0.1X diluted transfection method with a 60-minute incubation for transfection complex formation. 25% of the data were held back for validation of the trained models to prevent over-fitting. Training data included GFP-A, GFP-H, FSC-A, FSC-H, SSC-A and SSC-H values. All training parameters were retained at default settings. Results in Table 1 were obtained directly from the training app and the trained Fine Decision Tree model exported for use with additional datasets.

When applied to other datasets, a value for cell viability was obtained from the analysis by taking the confidence variable associated with each individual cell in the analysed population and averaging across the population.

### Flow cytometry

Flow cytometry experiments were performed using a Beckman Coulter Cytoflex S with standard methods. Gating strategies were specific to individual experiments and are illustrated in **Figure S1**. Prior to flow cytometry analysis, 50 µl PBS was added to each *Drosophila* cell culture in 96-well culture plates and cells were detached from the culture surface by manual pipetting. Flow cytometry data were analysed using CytExpert software (Beckman Coulter) and VDA scores calculated as described previously (25).

For analysis of VDA assays, GFP positive events were identified using a FITC gate similar to that shown in **Figure S1C**. However, this gate was applied to all events with no gating based on FSC or SSC. This is to allow inclusion of dying cells that still contain GFP signal, which contribute to changes in GFP distribution caused by cell viability phenotypes.

### VDA assays

Standard VDA assays were performed as described previously (25) with modifications to the protocol as described in the Results section and as illustrated in **Table 2**. All transfections were performed using Fugene HD transfection reagent (Promega).

**Table 2:** Contents of transfection mixtures for variants of the VDA method. All mixtures were generated in a total volume of 10 µl in PBS. Incubation times to allow transfection complex formation were 15 minutes unless otherwise stated in the results section. In particular, htVDA assays were performed with an incubation time of 60 minutes. All values for plasmids indicate ng per 10 µl transfection mix. Values for Fugene HD indicate µl per 10 µl transfection mix.

| Method | Gal4 | GFP | shRNA<br>(pValium20) | shRNA<br>(pELS) | Fugene HD |
| --- | --- | --- | --- | --- | --- |
| VDA | 90 | 20 | 90 |  | 0.6 |
| htVDA | 9 | 2 | 9 |  | 0.06 |
| htVDA |  | 2 |  | 18 | 0.06 |
| VDA-plus |  | 20 |  | 180 | 0.6 |
| Pooled VDA |  | 20 |  | 180 | 0.6 |

Proportions of reagents used in each transfection method are illustrated below.

Liquid handling automation was performed using a Mosquito LV Genomics liquid handling robot (SPT Labtech).

### Pooled VDA

Libraries for pooled VDA are described in **Table S1**. Transfections were performed as for VDA plus assays but using pooled libraries (KL1 to KL15) or sub-libraries instead of individual shRNA plasmids. Transfection mixtures were generated and then split prior to addition of half to S2R+ cells and half to NF1 deficient cells. Cytometry analysis was performed as described for VDA plus assays. Data were then analysed by calculating VDA scores for each library or sub-library in cell line and plotting results from each cell line against each other. For each library or sub-library, a ‘distance’ value was calculated, representing the orthogonal distance of each point on the scatter plot from a line indicating equal phenotype in both cell lines (**Figure S5**). This was defined as a straight line between the negative control point (transfected with *white* shRNA only) and 0,0.

## Competing Interest Statement

B.E.H. was a shareholder and founding director of Quest Genetics Ltd. between 2021 and March 2024. The remaining authors declare no competing interests.

## Data availability statement

All plasmids are available on request. Code for analysis of VDA data and trained machine learning models are available on request although we note that these must be edited or retrained for the specific cytometer used to generate the data to be analysed. All other data necessary for confirming the conclusions of the article are included within the article.

## Supplementary files

File S1: Supplementary results relating to Figure S2.

Figure S1: Illustration of example gating strategy for cytometry experiments

Figure S2: Optimisation of transfection conditions for VDA assays

Figure S3: Optimisation of miniaturised transfection method

Figure S4: Map of the pELS vector

Figure S5: Illustration of example data from the pooled VDA method

Table S1: Details of the pooled kinase shRNA library

## Supplementary figure legends

<u>Figure S2: Optimisation of transfection conditions for VDA assays</u>

**A-D)** Graphs indicate VDA signal from experiments performed with varying doses of Gal4 (A), shRNA (B) or both (D) or varying proportion of Gal4 and shRNA plasmid (C). Lines indicate average values from at least five replicate experiments and error bars indicate standard error of the mean. **E)** Graphs indicate relative cell viability measured using FSC/SSC gating from the same experiments described for panels A-C. **F)** Details of the panel are as for panel E but with varying proportion of Gal4 and shRNA in the transfections. **G)** Fold change in cell viability (measured using FSC/SSC gating) and transfection efficiency with and without recovery time between cell plating and transfection. Bars show the average of 30 replicate experiments and error bars indicate standard error of the mean. H-J) Lines indicate fold change in cell viability (measured using FSC/SSC gating; orange lines) and fold change in transfection efficiency (black lines) in transfections using varying volumes of transfection mixture. Lines show the average from five replicate experiments and error bars indicate standard error of the mean.

<u>Figure S3: Optimisation of miniaturised transfection method</u>

Lines indicate the % weight lost over time from at least four replicate experiments with either 5 µl PBS per well of a 384-well plate (red line) or 5 µl of PBS under 10 ul mineral oil per well of a 384-well plate (black line). Error bars indicate standard error of the mean.

## Acknowledgements

We would like to thank SPT Labtech for the generous loan of a Mosquito LV genomics liquid handling robot to allow initial testing of the htVDA method. This work was funded through Medical Research Council grant (MR/V009583/1) and a MRC Proximity to Development award.

