## Supplementary figures and images for "Enhanced methods for genetic assays in *Drosophila* cells"

### Supplementary Figure 1

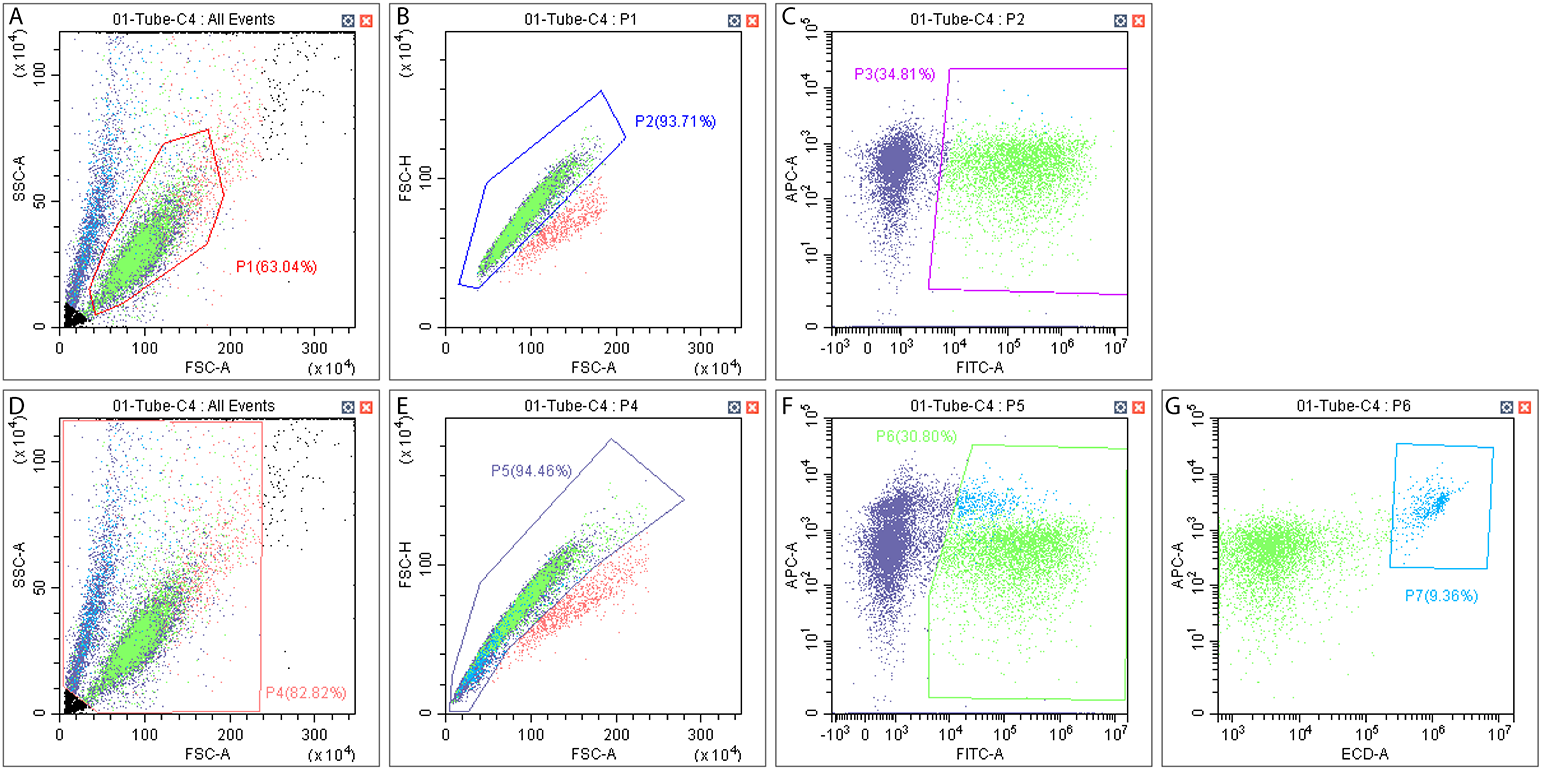

### Supplementary Figure 2

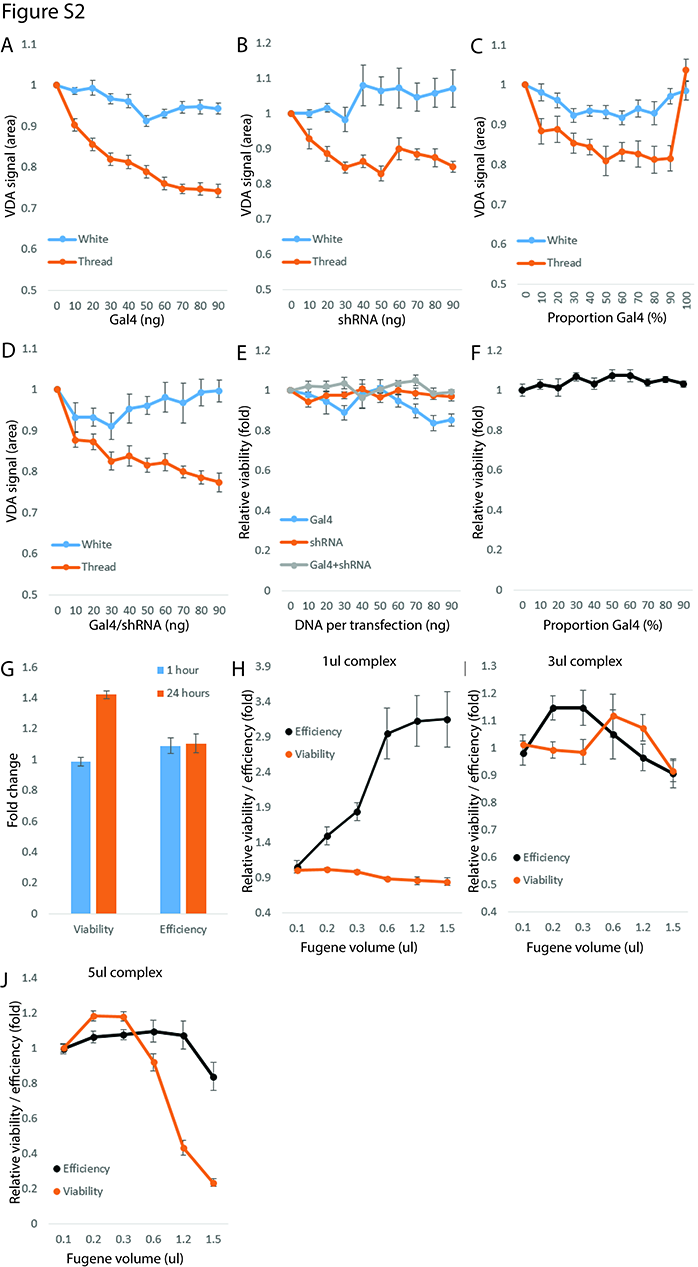

### Supplementary Figure 3

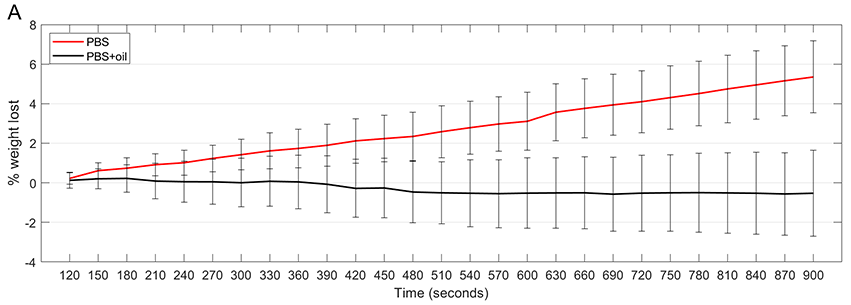

### Supplementary Figure 4

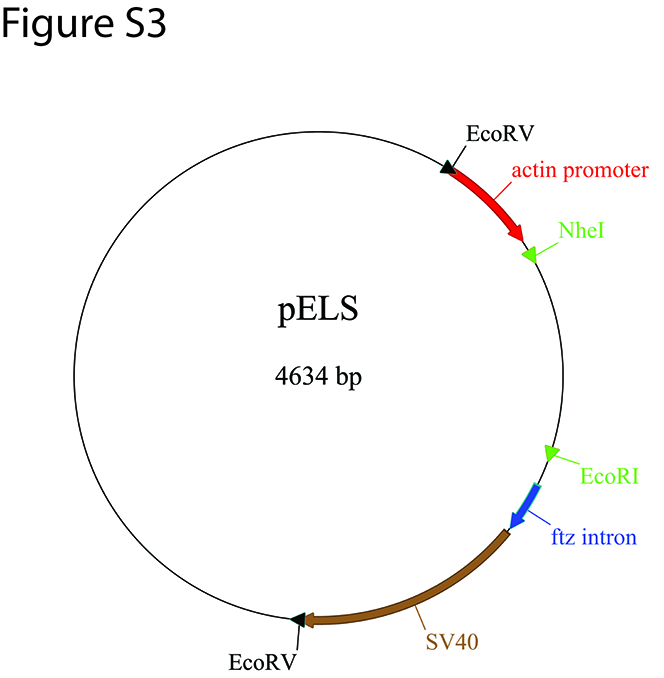

### Supplementary Figure 5

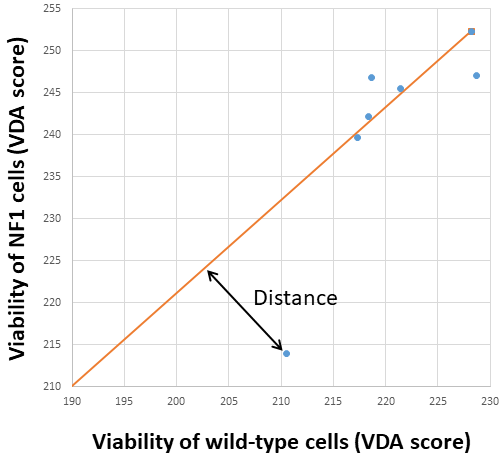
