## Supplementary File 1 for "Enhanced methods for genetic assays in *Drosophila* cells"

**Supplementary results:**

Optimisation of transfection conditions for VDA assays:

We sought to determine the most effective dose and ratio between the three plasmids to maximise VDA signal and maintain high transfection efficiency. We tested a range of transfection conditions, varying the total amount of Gal4 plasmid, shRNA plasmid or combined Gal4 and shRNA plasmid delivered as well as the ratio between the two plasmids (**Figure S2A-D**). For each condition, we measured the VDA signal in cells transfected with negative control (targeting the *white* gene) or positive control (targeting the *thread* gene to induce efficient cell death) shRNA reagents. We found that, VDA signal was stronger with higher doses of shRNA and Gal4 as indicated by the separation between the *white* and *thread* signals. However, despite this general trend, there existed a range of conditions over which VDA signals were broadly similar. For example, signals were relatively stable with Gal4 doses above 60 ng (**Figure S2A**), shRNA doses above 40 ng (**Figure S2B**) and Gal4:shRNA ratio between 1:1 and 9:1 (**Figure S2C**). By contrast, when the dose of Gal4 and shRNA were increased together, signal increased steadily up to the maximum dose (90 ng of each plasmid) (**Figure S2D**). These results are consistent with neither Gal4 nor shRNA plasmid being limiting across any condition tested. In addition, cell viability was relatively stable across all conditions tested for most plasmids (**Figure S2E-F**). One exception to this is that the Gal4 plasmid induced a reduction in viability at high dose (**Figure S2E**, blue line). This effect was also observed by a slight downward trend in the VDA signal of the negative control samples containing high Gal4 dose (**Figures S2A**, blue line).

Next, we went on to test the effect of additional transfection variables. First, we tested whether the cell seeding time influenced the toxicity and efficiency of transfection. Cells were transfected either immediately following seeding into a 96-well plate or 24-hours after seeding to allow recovery time. We found that the recovery time partially rescued the toxicity of transfection with no effect on transfection efficiency (**Figure S2G**), likely because stress due to the seeding process was resolved before the additional stress of transfection was applied.

Finally, we analysed the effects of varying the ratio of transfection reagent to DNA. This ratio was varied from 0.1 to 1.5 and was tested at three different total complex volumes. Interestingly, the effects of varying the ratio differed depending on the total complex volume. When 1 µl of complex was used, transfection efficiency increased and cell viability decreased as the ratio was increased (**Figure S2H**). However, when 3 µl or 5 µl of complex were used, peak efficiency and cell viability were achieved at intermediate ratios (**Figure S2H-J**). Nevertheless, a ratio of 3:1 of transfection reagent to DNA was approximately optimal in all cases.
